# Thermal cycling resets the irreversible liquid-to-solid transition of peptide condensates during aging

**DOI:** 10.1101/2025.04.16.649229

**Authors:** Abel Anwar, Tianchen Li, Yi Shen

**Affiliations:** School of Chemical and Biomolecular Engineering, The University of Sydney, PNR Building, Darlington NSW 2008, Australia

**Keywords:** Peptide self-assembly, Peptide condensates, Liquid-liquid phase separation, Liquid-to-solid transitions, Temperature Cycle

## Abstract

The ability of biomolecular condensates to reversibly dissolve and reform is crucial for maintaining cellular stability and functions. In metabolically active cells, stress granules can rapidly assemble and disassemble in response to environmental changes. However, as metabolic rates decline with aging, stress granules persist longer, disrupting mRNA translation and stress responses. Temperature, as a physical stimulus, plays a key role in controlling condensate formation, dissolution, and material properties. In this study, we explore how the reversibility of the liquid-to-solid transition of biomolecular condensates can be modulated by temperature change. Our findings reveal that aged condensates exhibit reduced responsiveness to external temperature stimuli. By using thermal cycling experiments to simulate repeated heat stress, we found that the time taken for irreversible fiber formation could be delayed up to 4.7-fold compared to condensates without thermal cycles. We also found the dissolution rate of condensates progressively slows as they age but remain more stable with thermal cycles. Importantly, our results indicate that continuous cycles of liquid-liquid phase separation and dissolution act as a reset mechanism, preserving the biomolecular condensates from further liquid-to-solid transition. These findings provide valuable insights into how aging impacts condensate behavior and highlight potential strategies to preserve cellular function through controlled phase transitions.

## 1. Introduction

Complex cellular systems are burdened with the important task of organizing and initiating necessary biochemical reactions to maintain homeostasis. A prominent idea involves partitioning intracellular components into distinct chemical environments via forming biomolecular condensates mediated by liquid-liquid phase separation (LLPS). This process plays an important role in stress mitigation mechanisms, controlling the formation of stress granules (SG). SGs are highly dynamic, liquid-like droplets important in regulating signaling pathways for mRNA translation formed transiently in the cytoplasm upon cellular stress across timescales of minutes to seconds. ^1–3^ However, this dynamic process slows down due to aging leading to the accretion of pathological fibrils. ^4, 5^ This may explain why certain neurodegenerative diseases such as ALS and FTLD disproportionately affect the elderly. ^6^ While such phase transition is at heart of both cellular functional and pathological activities, the mechanism of how it is regulated by external stimuli is not extensively explored, especially its correlation with aging time.

Biomolecular condensates, including SGs, can form under various forms of stress, including osmotic stress, heat shock, oxidative stress, nutrient starvation, and UV irradiation. ^7^ Of particular interest is the effect of heat stress on the assembly mechanism of condensates, given the ease with which it can be assessed *in vitro* compared to other stimuli. Temperature is a critical factor that directly modulates the LLPS of protein molecules. Specifically, the dual phase appears when the temperature is below the upper critical separation temperature (UCST) and/or above the lower critical separation temperature (LCST). Cells utilize such processes to detect temperature shock and maintain their fitness. ^8^ For example, Pab1 can form SGs under high temperatures to detect heat shock, ^9^ and Ddx4 can respond to cold shock and undergo LLPS. ^10^ It is also suggested that thermal sensing via protein condensation is conserved across multiple species and adaptive to specific thermal niches. ^11^ However, whether such thermal responsiveness is preserved over the condensate lifespan remains unclear.

In this study we utilize carboxybenzyl-protected diphenylalanine (z-FF) as a model peptide to examine the effect of temperature on condensate aging and LLPS dynamics. The FF repeat is a core recognition motif in the Aβ polypeptide implicated in Alzheimer’s disease. ^12^ Previous research has reported that z-FF can undergo LLPS through temperature control exhibiting UCST behavior. ^13–15^ Over a prolonged incubation period, the condensates solidify into fibrillar aggregates similar to pathological fibrils that can form in neurodegenerative diseases. These characteristics make this short peptide a suitable candidate for the study. We probed the thermoresponsivity of z-FF condensates via dissolution tests to assess the correlation between aging and impaired dynamics. To simulate cellular stress events, heat stress is applied by performing consecutive thermal cycles by using a microscope stage incubator. The impact of repetitive stress events on the condensate dynamics and eventual fibrillization is assessed. The results demonstrated the impact of repetitive thermal stress on LLPS and aging without changing any other conditions. They provide insights into the progressive decline of cellular metabolism in the elderly, particularly in the context of aging-related diseases and stress granule dynamics.

## 2. Materials and Methods

### 2.1. z-FF Liquid-liquid phase separation assay

Z-Phe-Phe-OH (z-FF) lyophilized powder was purchased from BACHEM, Switzerland. Dimethyl sulfoxide (analytical grade) was obtained from Ajax Finechem, Australia. Thioflavin T (ThT) was obtained from Sigma Aldrich, Australia.

A fresh stock of z-FF peptide solution was prepared by dissolving the lyophilized powder in DMSO and aliquoting it into centrifuge tubes. LLPS was then initiated by adding Milli-Q Ultrapure water, followed by gentle mixing before being pipetted into chamber wells for microscopy or stored in tubes for incubation. For microscopic imaging, a 200 µM stock solution of ThT in water was used to trigger the LLPS and stain the condensates or fibers. All microscopy images were captured using the confocal microscope (Nikon).

### 2.2. Phase diagrams

For the water-ratio phase diagram, the effect of the antisolvent was assessed by preparing z-FF final concentrations of 1, 5, 10, 15, 20, and 25 mg/ml across DMSO-water ratios of 1.5:1, 1.25:1, 1:1, 1:1.25, and 1:1.5 by volume in 500 µL aliquots. Samples were left to cool to room temperature for 30 mins before being imaged.

For the temperature-dependent phase diagram, similar z-FF final concentrations were prepared in a 1:1 DMSO-water ratio by volume in 600 µL aliquots. An identical set of samples was also prepared with ThT to a final concentration of 100 µM. Samples were incubated at 25, 35, 45, and 60 □in either an incubator (Cole-Parmer) or laboratory oven (Across International) for 2 hours and imaged at 10 min intervals to assess for turbidity and bulk gelation via tilting.

### 2.3. Confocal microscopic imaging

Microscope stage-top incubator samples were imaged using a Nikon confocal microscope and accompanying NIS-Elements software under a 20x objective and Nyquist optical zoom. Fluorescence due to the presence of ThT was measured via excitation at 488 nm and emission between 503 and 541 nm.

### 2.4. Turbidity assay

Fresh z-FF condensates were prepared in transparent 96-well plates with flat bottoms. When necessary, these condensates were further incubated at room temperature for aging. The turbidity of the peptide solution was measured using a microplate reader (CLARIOstar) in absorbance mode at 350 nm. ^16^ The plates were incubated inside the microplate reader for a temperature-dependent assay. The data was exported from MARS data analysis software and processed using MATLAB.

### 2.5. Incubator thermal cycling

Fresh z-FF condensates were prepared in a 1:1 by volume DMSO-water solution to a final concentration of 15 mg/mL in 600 µL aliquots. A thermal cycle was performed by heating the samples to 65 in a laboratory oven (Across International) for 5 mins, followed by ambient cooling to room temperature (25). Samples were then left at room temperature for 5, 20, 40, and 60 mins (aging period) before performing subsequent thermal cycles (48, 19, 11, and 7 cycles, respectively, across 8 hours). An additional two samples representing the non-cycled samples were prepared and subjected to constant incubation at 65 or 25. Images of the samples were taken at the end of each aging period to assess for turbidity and bulk gelation via tilting.

Characterization of z-FF morphology was performed at the equivalent total aging time of 240 min for each of the samples, corresponding to 48, 12, 6, and 4 thermal cycles for the 5, 20, 40, and 60 min aging periods, respectively. A small volume (112 µL) was pipetted from the centrifuge tubes containing the samples and stained with 8 µL of 1.5 mM ThT for microscopic imaging.

### 2.6. Microscopic thermal assay

For microscopic thermal cycling, a fresh z-FF in DMSO stock solution was gently mixed with a ThT-water solution in a 1:1 by volume ratio to a final z-FF concentration of 10 mg/mL in a centrifuge tube. An additional mixture was prepared without the peptide using identical volumes of DMSO and water to serve as a temperature reference. 200 µL of the sample and reference was transferred to an 18-well chambered cover glass (Cellvis) and sealed with parafilm. The chamber was placed inside the stage-top incubator (Tokai Hit) installed on the confocal microscope (Nikon) followed by immediately inserting the accompanying thermal probe into the reference mixture. Sample and incubator temperatures were controlled and measured via the STX-APP software and processed using MATLAB.

### 2.7. Stage-top incubator operation

Thermal cycling was achieved by manually operating the stage-top incubator where heating/cooling was managed by adjusting heater settings to the maximum/minimum. These settings are 65, 50, 50, and 45 for the top, stage, bath, and lens heater, respectively for the heating phase, and 10 □for all heaters for the cooling phase. The samples were heated to >45 □ to prompt dissolution, followed by ambient cooling for 15, 35, 65, 95 mins to achieve aging periods of 13, 30, 70, and 105 mins, respectively. After each aging period, the sample was heated to return to the dissolved state over 10 mins, or 18 mins for the 105 min aging period, and the cooling/heating cycle repeated until the formation of irreversible fibrils. Data was processed and plotted using MATLAB.

## 3. Results and Discussions

### 3.1. LLPS and Liquid-to-solid transitions of z-FF

z-FF undergoes LLPS when the good solvent (e.g. DMSO, ethanol, HFIP) is switched to an antisolvent such as water. ^17, 18^ A screening experiment is conducted at z-FF concentrations of 1, 5, 10, 15, 20, and 25 mg/ml under varying DMSO-water ratios of 1.5:1, 1.25:1, 1:1, 1:1.25, and 1:1.5 by volume where increasing water ratios are observed to be favorable for the triggering of LLPS (**Supporting information Figure S1**). To identify the aging progression of z-FF, a concentration of 15 mg/ml at a DMSO-water ratio of 1:1 by volume is selected, prepared in a centrifuge tube, and left to age at room temperature. The general progression of the LLPS and liquid-to-solid transition of z-FF is shown in (**Figure 1**). A white, turbid solution is initially observed which can flow smoothly when the tube is tilted provided the sample is relatively fresh. Upon further incubation for around 70 mins, a weak gel is formed which fails to flow as smoothly as a fresh sample when tilted. Microscopic images show that the gel is composed of a combination of stacked, aberrant-sized condensates as well as some fibrillar structures. Further aging (140 mins) results in a strong gel evidenced by a decrease in the turbidity (loss of dynamic condensates), followed by eventual bulk gelation at around 280 mins. The sample is composed predominantly of fibers or aged residues large enough to be observed as suspended structure within the tube.

**Figure 1.**
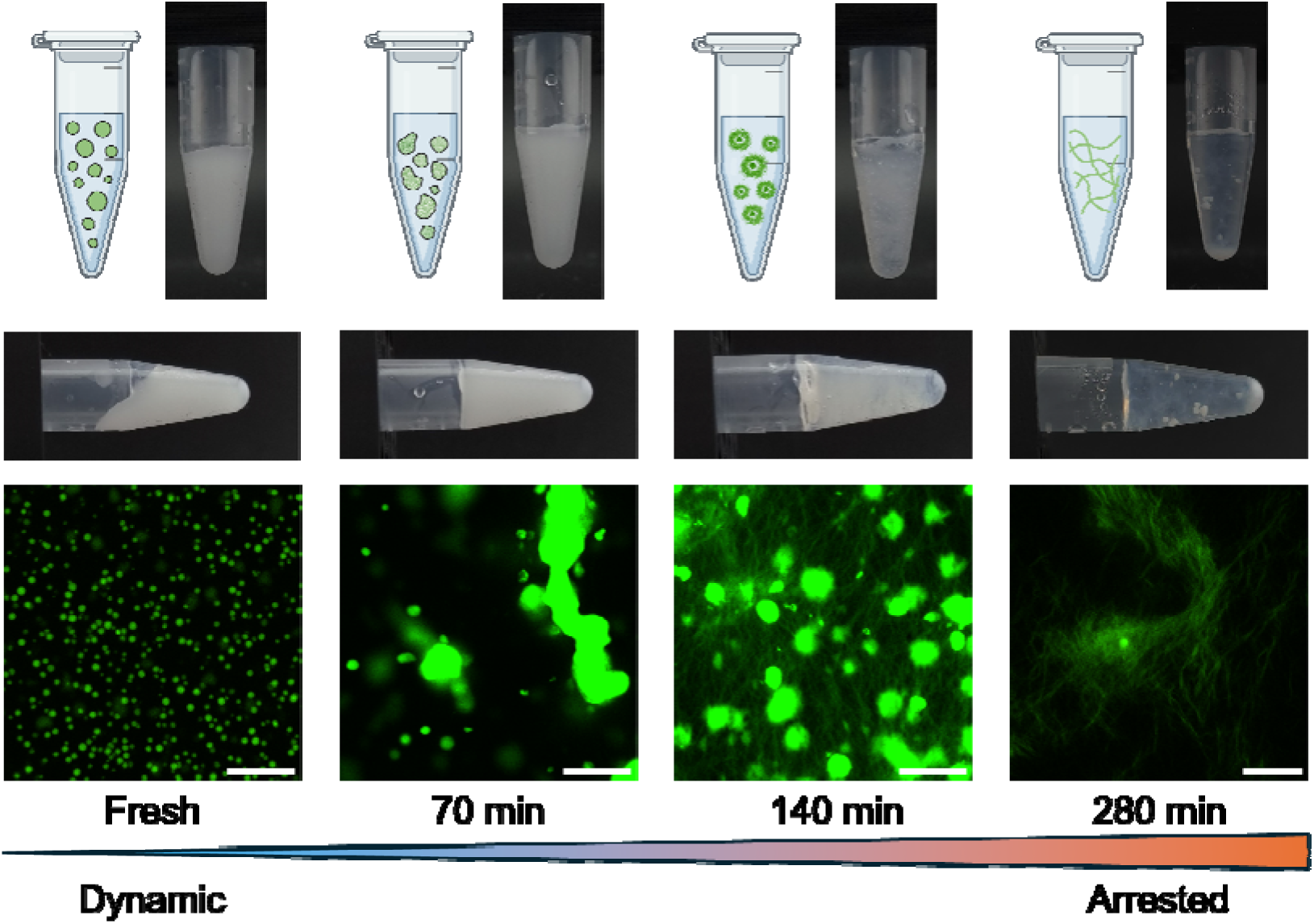
Different states of z-FF condensates at 15 mg/ml with a DMSO-H_2_O ratio of 1:1 by volume. Images from top to bottom are captured from centrifuge tubes, tilted centrifuge tubes, and confocal microscopy after staining, respectively (Scale bar: 20 µm). Total aging time from left to right: initial, 70 min, 140 min, and 280 min.

### 3.2. Temperature controls the LLPS and Liquid-to-solid transitions of z-FF

It has been reported that z-FF condensates display UCST behavior, and the critical temperature is concentration-dependent. ^15^ Here, besides the LLPS, we further investigate the aging of the z-FF condensate solution under different concentrations and temperatures within a two-hour span to map a preliminary phase diagram. z-FF is prepared at concentrations of 1, 5, 10, 15, 20, and 25 mg/ml in a DMSO-water ratio of 1:1 by volume and incubated at 25, 35, 45, and 60 □. An identical set of samples is also prepared with the addition of ThT to a final concentration of 100 µM. As described in the previous section, the aging of condensates can be roughly characterized by weak gel formation, which can be assessed by tilting the sample tubes and checking liquid mobility. The resultant concentration-temperature phase diagram is plotted at multiple time points, as shown in **Figure 2**. A similar phase diagram for the samples with ThT is provided in the supporting information (**Supporting Information Figure S2**). Weak gels are first observed in samples of high z-FF concentrations (20 – 25 mg/ml) and gradually spread to those with lower concentrations and temperatures. The results also show slower gelation dynamics under high temperatures, particularly when sample conditions are close to the boundary condition of LLPS. Interestingly, while the trend of lower temperatures favoring gelation is consistent across temperatures from 35 – 60 □, the gelation of the 25 samples seems to experience a slight delay until around 90 mins. Since the 25 – 35 □ temperature region exhibits identical states for each concentration, it is likely they have similar gelation dynamics that may be difficult to distinguish from each other due to the highly heterogeneous nature of z-FF gelation.

**Figure 2.**
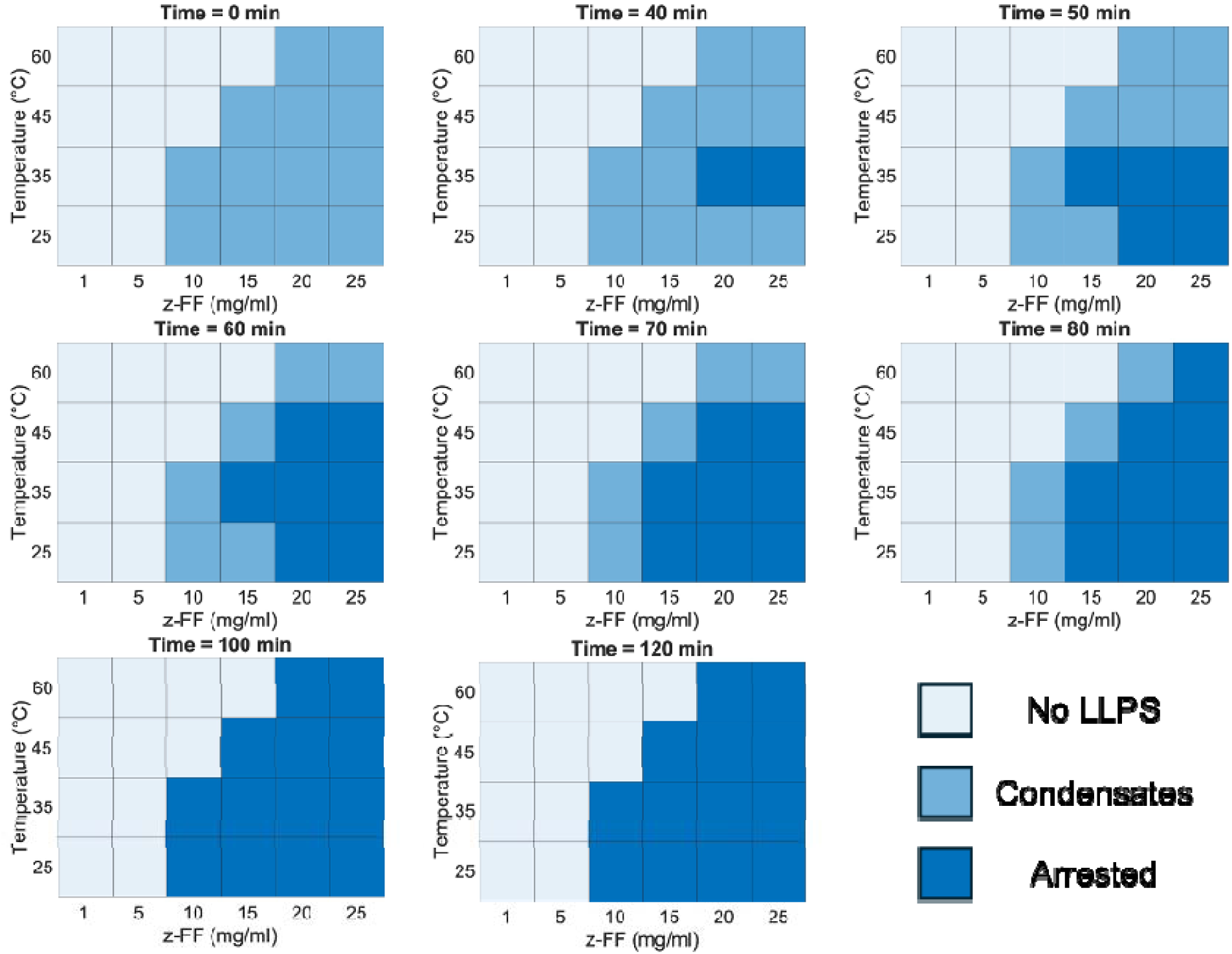
Phase diagrams of z-FF with DMSO-H_2_O ratio of 1:1 by volume at different time points without ThT.

Similar trends are observed in the phase diagram with ThT. However, there is an even greater distinction between the different incubation temperatures and the transition from liquid to solid. The presence of ThT has an inhibiting effect on the overall aging and gelation across all temperatures, with the first instance of weak gel formation occurring after 70 mins for the 15, 20, and 25 mg/ml samples at 25 □. Furthermore, high temperatures (60□) can completely preserve liquid mobility over 2 hours. The effect of ThT has been reported to increase gelation time by up to tenfold, as well as improve hydrogel rigidity for Fmoc-FF, a similar dipeptide. ^19^ The gelation mechanism is suggested as a two-step nucleation process, where the presence of ThT elongate the lag phase (condensate growth and protofibril nucleation) via disruption of the peptide packing interactions, resulting in an overall delay of the liquid-to-solid transition. ^19^ In both phase diagrams, the slowest aging dynamics are observed for conditions approaching the phase boundary corresponding to the metastable region between the binodal and spinodal curves. We hypothesize that repeatedly crossing the boundary may act to thermally reset LLPS (and subsequent liquid-to-solid transitions) and extend the lifespan of free z-FF capable of undergoing dynamic exchange between the dilute and concentrated phases.

### 3.3. Aged z-FF condensates display a slower responsiveness to the temperature

Next, we want to investigate whether the arrested condensates have a slower responsiveness to temperature change. The z-FF condensates (10 mg/ml) are incubated at 25 °C for different aging periods (0, 10, 50, 80 mins), followed by heating up to and incubated at 45 °C. When the incubator temperature is stabilized at 45 °C, the dissolution rate is significantly reduced for the aged condensates (**Figure 3a**). To achieve a more precise detection of condensate dissolution in response to temperature change, a similar experiment is done in the stage-top incubator on the microscope where the sample temperature instead of incubator temperature is measured. For this experiment, z-FF condensates at concentration 10 mg/ml with aging time of 13, 30, 70, and 105 mins incubated at 25 °C are prepared. The sample temperature is increased from 25 °C to 47 °C. It is interesting to observe that the 13 min- and 30 min-aged samples dissolve way before the sample temperature reaches 45 °C, whereas the dissolution of aged samples is more delayed (**Supporting information Figure S3**). To further visualize this, the normalized mean intensity reductions over temperature increments are calculated for each group (**Supporting Information, Figure S3**). As indicated in **Figure 3b**, the dissolution rate of 30 min-aged condensates is slightly lower than 13 min-aged condensates. The 70 and 105 min-aged condensates display a significantly lower dissolution rate than other groups. The reduced dissolution rate of aged condensates from both experiments may be due to the fact that the condensates are arrested with increased incubation time (Figure 2). Condensate arresting can significantly decrease the diffusivity over several degrees of magnitudes due to enhanced molecular interactions. The diffusive peptide flux towards the dilute phase is thus significantly decreased. ^20^

**Figure 3.**
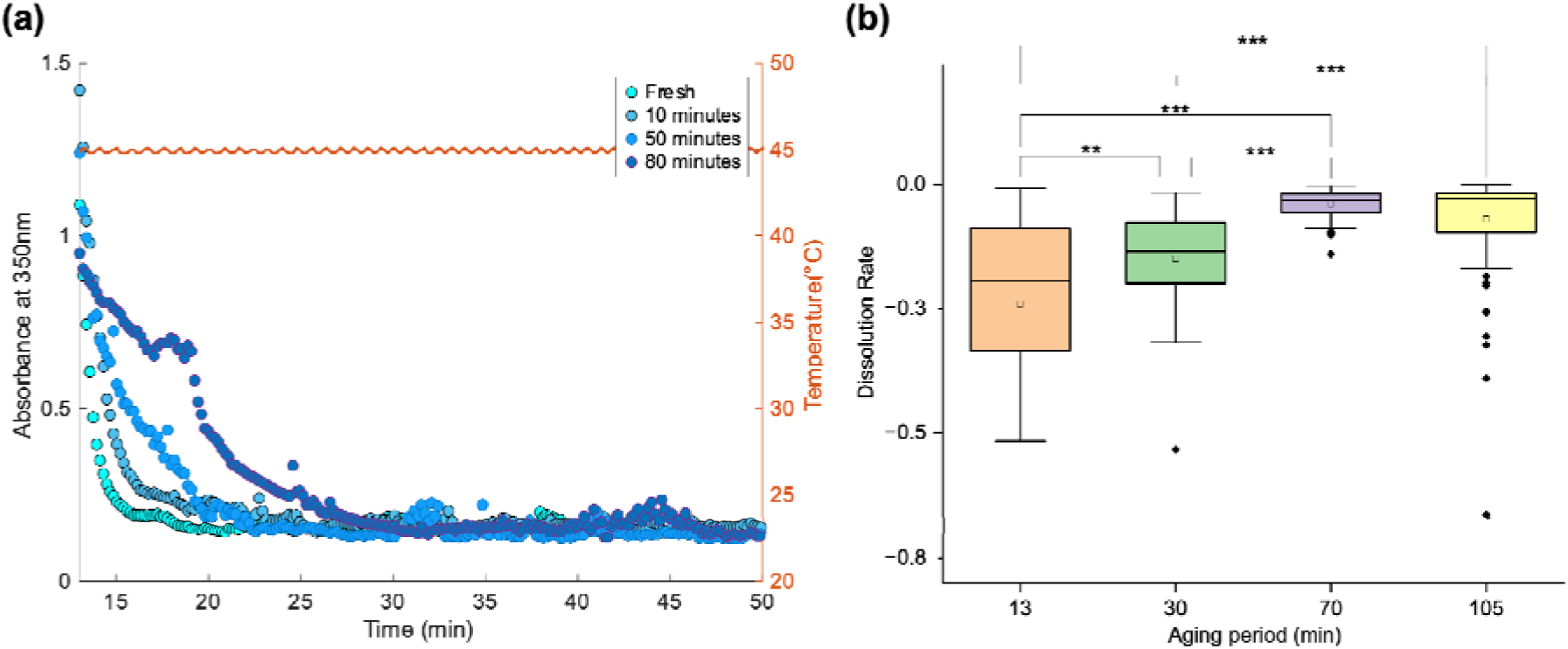
(a) Dissolution curve of different aged z-FF condensates. z-FF condensates pre-incubated for 0, 10, 50, and 80 mins at 25°C before heating up to 45°C. The turbidity is measured when the temperature is stabilized at 45°C for around 13 mins. (b) Dissolution rates of different aged condensates. z-FF condensates are pre-incubated for 13, 30, 70 and 105 min before heating up to 45°C. The dissolution rate is calculated by the step change of normalized intensity divided by the step change of temperature. (**Supporting information Figure S3**) * p<0.05, ** p<0.01, *** p<0.001.

### 3.4. Thermal cycling delays the aging of z-FF condensates

Following that, we want to examine whether continuous dissolution and reformation of condensates via thermal cycling can delay their aging. z-FF at 15 mg/ml with a 1:1 DMSO-water ratio is selected as a representative condition due to the faster aging dynamics indicated in the phase diagram (**Figure 2**). A control experiment representing the non-cycled condition is performed by incubation at 25 °C for 8 hours. The condensates form a weak gel at around 70 mins but retain their turbidity (**Figure 4b**). This indicates the coexistence of both aberrant condensates and solid fibers—the turbidity is implicit of light scattering associated with condensates, and the loss of sample mobility represents the critical point at which fiber growth has proliferated enough to support the sample volume. It is worth noting that the liquid-to-solid transition for z-FF so far has been observed to occur heterogeneously. That is, clusters of fibers form in random positions before extending out to nearby condensates, which ‘construct’ onto the fiber network. The initial formation of these fiber clusters may result from the radial fibril precursors. ^15^

**Figure 4.**
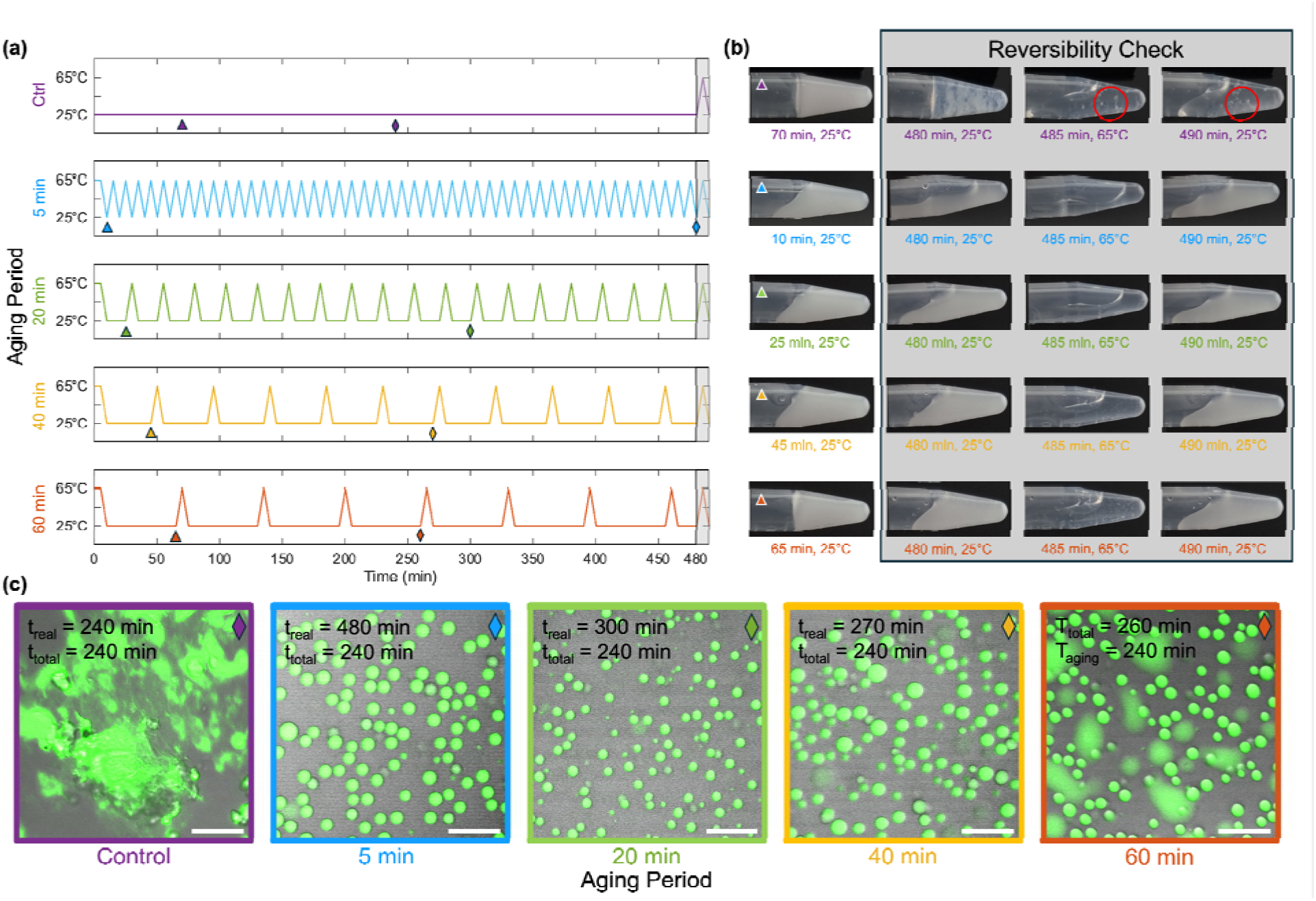
(a) Experimental design for the temperature cycling assay in centrifuge tubes. (b) Representative images of each testing group at different time points. The time points when the images are taken are marked as triangles in (a). (c) Microscope images of ThT-stained sample from all groups with the same total aging time of 240 mins (marked as diamonds in (a)).

Further incubation (140 mins) results in a decrease in turbidity and the formation of dense fiber networks before becoming mostly transparent by 280 mins with significant visible fiber structures (**Supporting Information Figure S4**). At 240 mins of aging time, significant aggregates and fiber structures are observed, indicating that most of the z-FF molecules have already undergone the irreversible liquid-to-solid transition (**Figure 4c**). By 8 hours of incubation, a transparent gel with a suspended fibrillar network is achieved. When the gel is further heated to 65 °C, it recovers as a transparent liquid with notable residues left behind (**Figure 4b**). The reduced turbidity can be associated with the dissolution of condensates embedded or wetted on the fibrillar network. When the condensates are dissolved, the fibrillar network also disassociates and fails to support the hydrogel, leading to recovery of mobility. Interestingly, when the temperature is restored to 25 °C, the turbidity of the solution shows negligible recovery (**Supporting Video 1**). We speculate as most z-FF molecules have served as building blocks of solid aggregates, the remaining z-FF in the solution cannot form large condensates.

To probe the effect of thermal cycling on condensate aging, four identical samples of z-FF (15 mg/ml) are subjected to thermal cycling via rapid heating to 65□ for 5 mins, followed by cooling to 25 °C. The samples are incubated periodically at 25 °C for 5, 20, 40, and 60 min aging periods before performing subsequent cycles as indicated in **Figure 4a**. Given the varying aging periods for each sample, the total aging time is considered as the sum of the aging periods. In contrast, the experimental (or ‘real’) time considers the entire length of the dissolution-reformation cycles. (**Supporting information Figure S4**)

All samples maintain turbidity even after up to 8 hours of continuous dissolution-reformation cycles (**Supporting Information Figure S4**). The sample with the 60 min aging period expectedly shows the most resemblance to the non-cycled control with sample immobilization and formation of a weak gel. However, this also shows some variance between cycles with liquid-like mobility observed on the 2^nd^, 3^rd^, and 5^th^ cycle (**Supporting Information Figure S5**). The 60 min aging period is only 10 mins shorter than the expected time required for initial weak gel formation whilst also showing varying degrees of immobilization (a result of the heterogenous liquid-to-solid transition). This corroborates the hypothesis that thermal cycling acts, to some degree, as a ‘reset’ mechanism. Furthermore, the transparent gel formation is never observed for the cycled samples well after the expected total aging time of approximately 140 mins. At 8 hours of real time following a final dissolution, solid debris is apparent in the 60 and 40 min aging period samples (total aging time of 440 and 425 mins, respectively) (**Figure 4b**), indicating that although the liquid-to-solid transitions of z-FF cannot be prevented over these timespans, they can be delayed.

Each sample is stained with ThT at an equivalent total aging time of 240 mins and imaged via confocal microscopy, yielding highly spherical condensates and no observable fibers. This is in stark contrast to the non-cycled control where upon staining no condensates are observed at all. A trend is apparent with an increase in aberrant structures (non-spherical condensates) with longer aging periods (**Figure 4c**), supporting the idea that longer aging periods are more likely to be further along in the liquid-to-solid progression. This would suggest that shorter thermal cycle may be beneficial to delaying condensate solidification and fibrillization.

To assess the impact of aging on condensate dynamics and recovery ability, we probe the microscale activity of the z-FF condensates during thermal cycles using confocal microscopy and a stage-top incubator to simultaneously allow for modulation of the sample temperature and imaging (**Figure 5a**, see also **Materials and Methods**). The operating temperature limit of the stage-top incubator is 50□, which is notably cooler than the oven heating temperature of 65 used in the previous macroscale cycling experiments. Therefore, a lower concentration of z-FF at 10 mg/ml is used in this study due to exhibiting a UCST of around 34 – 35□. A concentrated peptide stock is mixed with a water-ThT solution (200 µM) to a final concentration of 10 mg/ml with a DMSO-water ratio of 1:1 by volume. Note that although ThT is known to exhibit increases in fluorescence by several orders of magnitude upon binding to the beta-sheet structure of amyloid fibrils ^21^, ThT can also be effectively recruited to condensates through aromatic interactions and shows a fluorescent signal due to its sensitivity to viscosity. ^22^ Thus, using ThT to check the formation of beta-sheet structure inside the condensates may not be precise. Nevertheless, ThT will display a negligible signal if the condensates are fully dissolved and a notable signal if there are insoluble aggregates after dissolution. In this case, we can still judge whether the liquid-to-solid transitions occur.

**Figure 5.**
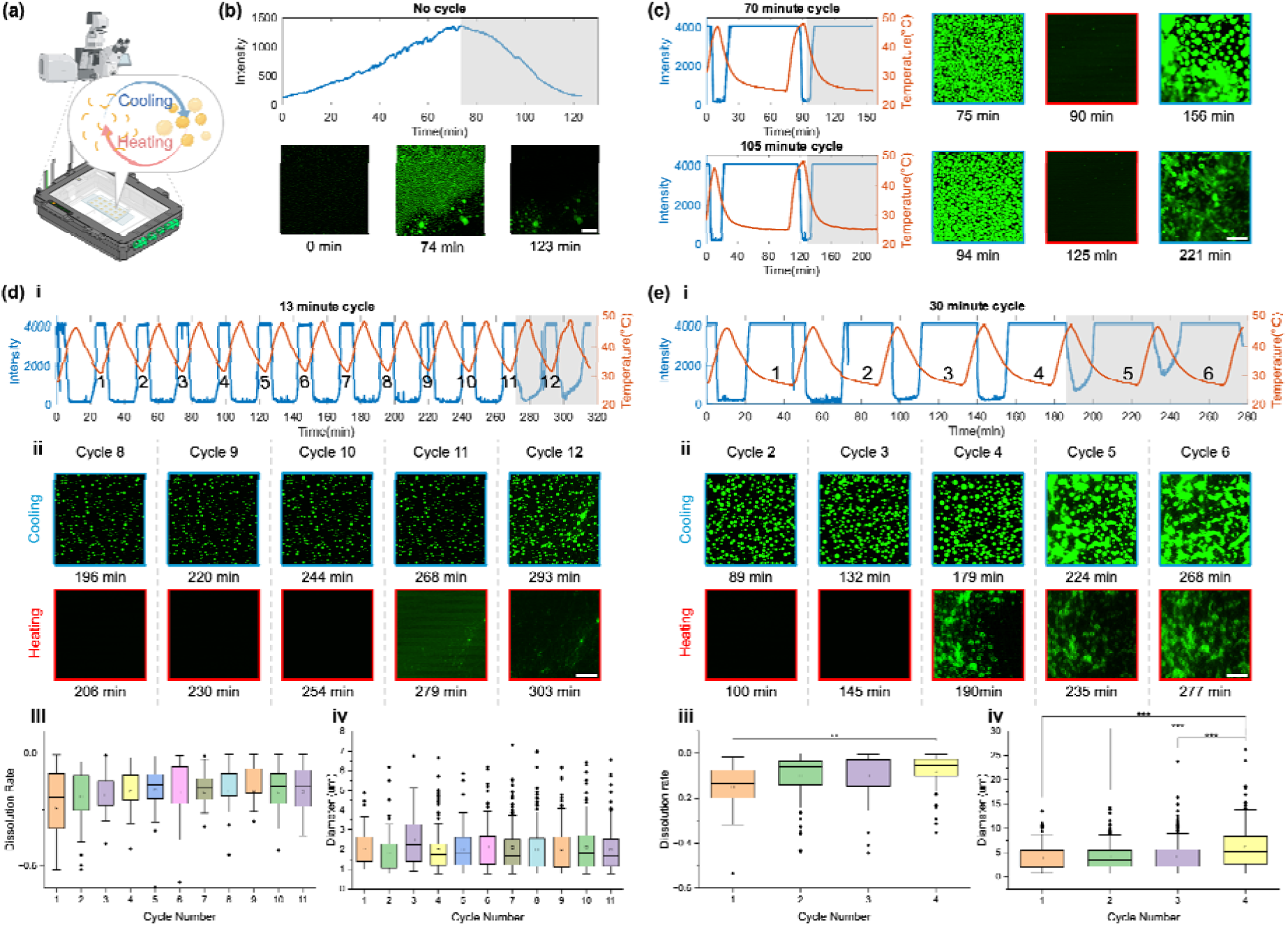
(a) Experimental setup. Peptide condensates are incubated in a stage-top incubator, settling on the confocal microscope. (b) Prolonged incubation of z-FF condensates (10 mg/ml) under 25 ℃ leads to aging and aggregation. Top: Mean intensity of the observed region over time. Bottom: Representative fluorescent images of the condensates, arrested condensates, and aggregates. Scale bar: 50 µm. (c) 70 and 105 min aging periods fail to fully dissolve the condensates after the first cycle. Left: Max fluorescent intensity and temperature profile. Right: Representative microscopy images in the second cycle. Scale bar: 20 µm. (d, e) z-FF condensates undergo dissolution-reformation cycles with an aging period of (d) 13 mins and (e) 30 mins. (i) Maximum intensity and temperature profile over time. The sudden decrease in intensity at the low-temperature region is due to microscope refocusing. The regions where condensate aging and aggregation are observed are highlighted in grey. (ii) Representative microscopy image during cycles, with corresponding time points and temperature labelled at the bottom. Scale bar: 20 µm. (iii) Dissolution rates calculated by the step change of mean intensity over the step change of temperature among all cycles (**Supporting information Figure S6**). (iv) Size distribution of condensates among cycles. * p<0.05, ** p<0.01, *** p<0.001.

A control experiment is conducted by incubating at 25 over 120 mins, which is the expected gelation time based on previous phase diagrams (**Supporting Information Figure S2**). An increase in the mean fluorescence intensity is associated with the Ostwald ripening of condensates until around 70 mins when extensive aberrant-sized condensates are observed. These large condensates of abnormal shape indicate the early fiber network which tends to form radially about these condensates. They can even be seen forming as early as 45 mins within a small region locally followed by forming an extended fibrillar network around 60-70 mins (**Figure 5b**). This aligns with previous observations of heterogeneous aging phenomena where the initial radial cluster can form as early as 30–45 mins. Despite the presence of these initial fiber networks, they are too sparse to support the entire sample volume and hence liquid mobility is still preserved. The fiber network continues to proliferate by seizing nearby condensates to construct onto the network, forming thin fibers and decreasing the fluorescence intensity (**Supporting Video 2**). At the same time, condensates appear to dissolve due to a decrease in local concentration caused by fiber formation. This releases the ThT back into the environment and thus decreases fluorescence intensity. By around 123 mins, the fluorescence intensity is minimal and contributed to only by the remaining fiber network, which agrees with the gel formed by 120 mins indicated by the phase diagram (**Supporting information Figure S2**). In this regard, the incubator phase diagrams can be seen as representative of the bulk sample gelation or dynamics, while the stage-top incubator microscopy can provide detail regarding sample aging kinetics and initial fiber formation.

Thermal cycling is also performed on samples of identical composition to the 25□ control by varying the time permitted for the z-FF to exist in the phase-separated state (aging period). These periods are 13, 30, 70, and 105 mins long, achieved by cooling the sample for 15, 35, 65, and 95 mins, respectively, followed by heating for 10 mins (**Figure** 5**c-e****, Supporting Video 3-6**). Sample heating results in peak temperatures of 46 – 48□, which is well above the observed critical dissolution temperature of approximately 34 – 35□, ensuring that the z-FF condensates are adequately dissolved. Sample max intensity is recorded and shows periodic peaks and troughs inverse to those in the temperature profile, corresponding to the phase-separated and dissolved states at low and high temperatures. While fiber formation could not be prevented, they could be significantly delayed by up to 4.7 times the total aging time of the non-cycled control. The effect of prolonging the ‘lifespan’ of the condensate phase is most apparent in the shortest aging periods (30 and 13 min) where fiber formation is observed on the 4^th^ and 11^th^ cycles, respectively (**Figure 5d, e**). These correspond to total aging times of approximately 120 and 143 mins, respectively, a roughly 4 to 4.7-fold increase in condensate lifespan over the control. This contrasts with the longer aging periods (70 and 105 min), which can only undergo one thermal cycle before fibrilization (**Figure 5c**). Therefore, an intrinsic link exists between the aging period and number of possible thermal cycles. Shorter aging periods directly result from more frequent thermal cycles for a fixed total aging time. This leads to preserving the free peptide over time, delaying the eventual liquid-to-solid transition.

The dissolution rates and size distribution of condensates within the 13 min and 30 min cycles are measured. No significant change among the dissolution rates and condensate size distributions before aggregate formation for the 13 min aging period is observed (**Figure 5e, iii, iv**). For the 30 min aging period, there is a slight reduction in dissolution rate on the 4^th^ cycle (**Figure 5e, iii**). A growth of condensate size can also be observed in the 4^th^ cycle before the aggregation is detected (**Figure 5e, iv**). When the slightly aged condensates are dissolved, they may not fully convert back to the free peptide monomers. Instead, the nanoclusters or nanofibril nuclei may remain and accumulate over time. This potentially decreases the energy barrier for the LLPS and condensate ripening when the temperature is decreased in the next cycle, leading to the formation of larger condensates followed by liquid-to-solid transitions. It is previously reported that during the light-induced cycles of p62 protein condensates, PolyQ aggregates can be gradually accumulated on the interface of the p62 condensates. ^23^ We speculate that the z-FF condensates undergo a similar surface-mediated aging progress, indicated by the core-shell structure left over after the dissolution of aged condensates (**Figure 5e, ii**). By contrast, shorter cycle periods can potentially hinder the formation and accumulation of such nanoclusters, thus delaying the liquid-to-solid transition.

It is important to note that these observations represent the early stages of the sample liquid-to-solid transition (i.e., corresponding to samples with liquid-like mobility or very weak turbid gels) rather than the final, gelled state (transparent). This is evidenced by the presence of dynamic condensates in other positions within the sample and the ability of the condensates to nucleate onto the fibers upon cooling which results in the recovery of turbidity. In the context of biological systems where both the phase-separated and dispersed states of proteins are relevant, the ‘real’ time also becomes relevant as the age of the organism itself. Then, the protein’s lifespan should include the length of time kept in both the dispersed and phase-separated state. By these definitions, the effect of thermal cycling becomes even more apparent. For the control experimental, the real-time is the same as the total aging time and takes approximately 30 mins for initial fiber formation or 70 mins for weak bulk gelation. This is significantly shorter than the real time for the 13 and 30 min aging periods which are at minimum 268 and 179 mins, respectively. Nevertheless, regardless of the definition of time (total aging time or experimental time), the result of prolonging condensate lifespans via thermal cycling is consistent across both comparisons.

## 4. Conclusions

In this study, we utilize a novel approach involving a microscopic stage-top incubator to probe the effects on condensate dynamics and thermoresponsivity of a model dipeptide system by simulating repeated heat stress events. We report that the z-FF dipeptide, which readily undergoes LLPS under solvent-exchange with UCST behavior, exhibits inhibited dynamics and thermoresponsivity under prolonged aging. However, by performing repeated rapid dissolution-reformation cycles (thermal cycles), the dynamic nature of the condensates in terms of lifespan can be preserved by up to 4.7-fold in the best case, and the eventual liquid-to-solid transition to a fibrillar hydrogel network is delayed. We propose that continuously cycling z-FF between the phase-separated and dispersed state via temperature control is a form of thermal reset that can preserve the free z-FF in the system. Furthermore, a link between aging and the number of possible cycles before irreversible fibrillization is also observed with more frequent cycles, akin to regular metabolism, being favorable for preserving free z-FF.

While our study reveals the link between aging and condensate stress response for a simplistic dipeptide system, potential future studies may extend into using more physiologically relevant proteins to ascertain if the reported behavior applies more generally. It should be noted that although critical temperatures for the studied z-FF concentrations are close to the physiological temperature range of 36-37 ℃, experiments involving thermal cycling are conducted over a significantly wider range (25–60 ℃), indicating limited practicality in biological systems or in vivo studies. However, as mentioned previously, environmental stress is not exclusive to thermal stress. Other stress-induced cycling methods (osmotic, pH, oxidative, light, etc.) could reasonably be employed to simulate repeated stress events. ^24, 25^ Although whether these methods can achieve a similar ability to preserve free protein remains unclear, they may provide insight towards or inspire potential therapeutic strategies to preserve cellular function.

## ASSOCIATED CONTENT

### Supporting Information

The following files are available free of charge.

Supporting information.pdf

Supporting video 1 – Reversibility_Check.mp4 - A final reversibility check for z-FF condensates with different aging periods as indicated in Figure 4b.

Supporting video 2 – zFF_aging.mp4 – Aging of z-FF condensates. Clusters of fibers form in random positions. The fiber network continues to proliferate by seizing nearby condensates to construct onto the network, forming thin fibers and decreasing the fluorescence intensity.

Supporting video 3 – zFF10_13min_cycle.avi - Temperature cycle of z-FF at 10 mg/ml with an aging period of 13 mins.

Supporting video 4 – zFF10_30min_cycle.avi - Temperature cycle of z-FF at 10 mg/ml with an aging period of 30 mins.

Supporting video 5 – zFF10_70min_cycle.avi - Temperature cycle of z-FF at 10 mg/ml with an aging period of 70 mins.

Supporting video 6 – zFF10_105min_cycle.avi - Temperature cycle of z-FF at 10 mg/ml with an aging period of 105 mins.

## Author Contributions

The manuscript was written through contributions of all authors. All authors have given approval to the final version of the manuscript. † These authors contributed equally.

## ABBREVIATIONS

z-FF: carboxybenzyl-protected diphenylalanine
LLPS: liquid-liquid phase separation
UCST/LCST: upper/lower critical separation temperature
ThT: Thioflavin T
HFIP: 1,1,1,3,3,3-hexafluoro-2-propanol
SG: stress granules

## References

(1) Buchan, J. R.; Parker, R. Eukaryotic Stress Granules: The Ins and Outs of Translation. Molecular Cell 2009, 36 (6), 932–941. DOI: 10.1016/j.molcel.2009.11.020.

(2) Hyman, A. A.; Weber, C. A.; Jüelicher, F. Liquid-Liquid Phase Separation in Biology. Annu Rev Cell Dev Bi 2014, 30, 39–58. DOI: 10.1146/annurev-cellbio-100913-013325.

(3) Wippich, F.; Bodenmiller, B.; Trajkovska, Maria G.; Wanka, S.; Aebersold, R.; Pelkmans, L. Dual Specificity Kinase DYRK3 Couples Stress Granule Condensation/Dissolution to mTORC1 Signaling. Cell 2013, 152 (4), 791–805. DOI: 10.1016/j.cell.2013.01.033.

(4) Kim, H. J.; Kim, N. C.; Wang, Y.-D.; Scarborough, E. A.; Moore, J.; Diaz, Z.; MacLea, K. S.; Freibaum, B.; Li, S.; Molliex, A.;, et al. Mutations in prion-like domains in hnRNPA2B1 and hnRNPA1 cause multisystem proteinopathy and ALS. Nature 2013, 495 (7442), 467–473. DOI: 10.1038/nature11922.

(5) Molliex, A.; Temirov, J.; Lee, J.; Coughlin, M.; Kanagaraj, A. P.; Kim, H. J.; Mittag, T.; Taylor, J. P. Phase separation by low complexity domains promotes stress granule assembly and drives pathological fibrillization. Cell 2015, 163 (1), 123–133. DOI: 10.1016/j.cell.2015.09.015 From NLM.

(6) Niccoli, T.; Partridge, L.; Isaacs, A. M. Ageing as a risk factor for ALS/FTD. Human Molecular Genetics 2017, 26 (R2), R105–R113. DOI: 10.1093/hmg/ddx247 (acccessed 3/22/2025).

(7) Cao, X.; Jin, X.; Liu, B. The involvement of stress granules in aging and aging-associated diseases. Aging Cell 2020, 19 (4), e13136. DOI: 10.1111/acel.13136.

(8) Watson, J. L.; Seinkmane, E.; Styles, C. T.; Mihut, A.; Krüger, L. K.; McNally, K. E.; Planelles-Herrero, V. J.; Dudek, M.; McCall, P. M.; Barbiero, S.;, et al. Macromolecular condensation buffers intracellular water potential. Nature 2023, 623 (7988), 842–852. DOI: 10.1038/s41586-023-06626-z.

(9) Saad, S.; Cereghetti, G.; Feng, Y.; Picotti, P.; Peter, M.; Dechant, R. Reversible protein aggregation is a protective mechanism to ensure cell cycle restart after stress. Nature Cell Biology 2017, 19 (10), 1202–1213. DOI: 10.1038/ncb3600.

(10) Nott, Timothy J.; Petsalaki, E.; Farber, P.; Jervis, D.; Fussner, E.; Plochowietz, A.; Craggs, T. D.; Bazett-Jones, David P.; Pawson, T.; Forman-Kay, Julie D.; et al. Phase Transition of a Disordered Nuage Protein Generates Environmentally Responsive Membraneless Organelles. Molecular Cell 2015, 57 (5), 936–947. DOI: 10.1016/j.molcel.2015.01.013.

(11) Keyport Kik, S.; Christopher, D.; Glauninger, H.; Hickernell, C. W.; Bard, J. A. M.; Lin, K. M.; Squires, A. H.; Ford, M.; Sosnick, T. R.; Drummond, D. A. An adaptive biomolecular condensation response is conserved across environmentally divergent species. Nature Communications 2024, 15 (1), 3127. DOI: 10.1038/s41467-024-47355-9.

(12) Reches, M.; Gazit, E. Casting metal nanowires within discrete self-assembled peptide nanotubes. Science 2003, 300 (5619), 625–627. DOI: 10.1126/science.1082387.

(13) Yuan, C.; Levin, A.; Chen, W.; Xing, R.; Zou, Q.; Herling, T. W.; Challa, P. K.; Knowles, T. P. J.; Yan, X. Nucleation and Growth of Amino Acid and Peptide Supramolecular Polymers through Liquid–Liquid Phase Separation. Angewandte Chemie International Edition 2019, 58 (50), 18116–18123. DOI: 10.1002/anie.201911782.

(14) Shen, Y.; Ruggeri, F. S.; Vigolo, D.; Kamada, A.; Qamar, S.; Levin, A.; Iserman, C.; Alberti, S.; George-Hyslop, P. S.; Knowles, T. P. J. Biomolecular condensates undergo a generic shear-mediated liquid-to-solid transition. Nat Nanotechnol 2020, 15 (10), 841–847. DOI: 10.1038/s41565-020-0731-4.

(15) Zhou, P.; Xing, R.; Li, Q.; Li, J.; Yuan, C.; Yan, X. Steering phase-separated droplets to control fibrillar network evolution of supramolecular peptide hydrogels. Matter 2023, 6 (6), 1945–1963. DOI: 10.1016/j.matt.2023.03.029.

(16) Roberts, S.; Harmon, T. S.; Schaal, J. L.; Miao, V.; Li, K.; Hunt, A.; Wen, Y.; Oas, T. G.; Collier, J. H.; Pappu, R. V.;, et al. Injectable tissue integrating networks from recombinant polypeptides with tunable order. Nature Materials 2018, 17 (12), 1154–1163. DOI: 10.1038/s41563-018-0182-6.

(17) Brown, N.; Lei, J.; Zhan, C.; Shimon, L. J. W.; Adler-Abramovich, L.; Wei, G.; Gazit, E. Structural Polymorphism in a Self-Assembled Tri-Aromatic Peptide System. ACS Nano 2018, 12 (4), 3253–3262. DOI: 10.1021/acsnano.7b07723.

(18) Yuan, C.; Levin, A.; Chen, W.; Xing, R.; Zou, Q.; Herling, T. W.; Challa, P. K.; Knowles, T. P. J.; Yan, X. Nucleation and Growth of Amino Acid and Peptide Supramolecular Polymers through Liquid–Liquid Phase Separation. Angewandte Chemie 2019, 131 (50), 18284–18291. DOI: 10.1002/ange.201911782.

(19) Tikhonova, T. N.; Rovnyagina, N. N.; Arnon, Z. A.; Yakimov, B. P.; Efremov, Y. M.; Cohen-Gerassi, D.; Halperin-Sternfeld, M.; Kosheleva, N. V.; Drachev, V. P.; Svistunov, A. A.;, et al. Mechanical Enhancement and Kinetics Regulation of Fmoc-Diphenylalanine Hydrogels by Thioflavin T. Angewandte Chemie International Edition 2021, 60 (48), 25339–25345, 10.1002/anie.202107063. DOI: 10.1002/anie.202107063 (acccessed 2022/10/23).

(20) Jawerth, L.; Fischer-Friedrich, E.; Saha, S.; Wang, J.; Franzmann, T.; Zhang, X.; Sachweh, J.; Ruer, M.; Ijavi, M.; Saha, S.;, et al. Protein condensates as aging Maxwell fluids. Science 2020, 370 (6522), 1317–1323. DOI: doi:10.1126/science.aaw4951.

(21) Biancalana, M.; Koide, S. Molecular mechanism of Thioflavin-T binding to amyloid fibrils. Biochimica et Biophysica Acta (BBA) - Proteins and Proteomics 2010, 1804 (7), 1405–1412. DOI: 10.1016/j.bbapap.2010.04.001.

(22) Leppert, A.; Feng, J.; Railaite, V.; Bohn Pessatti, T.; Cerrato, C. P.; Mörman, C.; Osterholz, H.; Lane, D. P.; Maia, F. R. N. C.; Linder, M. B.;, et al. Controlling Drug Partitioning in Individual Protein Condensates through Laser-Induced Microscale Phase Transitions. Journal of the American Chemical Society 2024, 146 (28), 19555–19565. DOI: 10.1021/jacs.4c06688.

(23) Choi, C. H.; Lee, D. S. W.; Sanders, D. W.; Brangwynne, C. P. Condensate interfaces can accelerate protein aggregation. Biophys J 2024, 123 (11), 1404–1413. DOI: 10.1016/j.bpj.2023.10.009 From NLM.

(24) Li, T.; Ilhamsyah, D.; Tai, B.; Shen, Y. Functional Biomaterials Derived from Protein Liquid–Liquid Phase Separation and Liquid-to-Solid Transition. Advanced Materials n/a (n/a), 2414703. DOI: 10.1002/adma.202414703.

(25) Pattanayak, G. K.; Liao, Y.; Wallace, E. W. J.; Budnik, B.; Drummond, D. A.; Rust, M. J. Daily Cycles of Reversible Protein Condensation in Cyanobacteria. Cell Reports 2020, 32 (7), 108032. DOI: 10.1016/j.celrep.2020.108032.

