## Supporting Information for "Thermal cycling resets the irreversible liquid-to-solid transition of peptide condensates during aging"

The following documents are provided free of charge:

Supporting information.pdf
Supporting video 1 – Reversibility_Check.mp4 - A final reversibility check for z-FF condensates with different aging periods as indicated in Figure 4b.
Supporting video 2 – zFF_aging.mp4 – Aging of z-FF condensates. Clusters of fibers form in random positions. The fiber network continues to proliferate by seizing nearby condensates to construct onto the network, forming thin fibers and decreasing the fluorescence intensity.
Supporting video 3 – zFF10_13min_cycle.avi - Temperature cycle of z-FF at 10 mg/ml with an aging period of 13 mins.
Supporting video 4 – zFF10_30min_cycle.avi - Temperature cycle of z-FF at 10 mg/ml with an aging period of 30 mins.
Supporting video 5 – zFF10_70min_cycle.avi - Temperature cycle of z-FF at 10 mg/ml with an aging period of 70 mins.
Supporting video 6 – zFF10_105min_cycle.avi - Temperature cycle of z-FF at 10 mg/ml with an aging period of 105 mins.

The supporting movies exceed the file size limit of bioRxiv. The original movies can be found at this URL:

<https://unisydneyedu-my.sharepoint.com/:f:/g/personal/aanw5541_uni_sydney_edu_au/EhjxgvLL0hJJlIomcbt0zuQBBZ2UUE-4pb5_RdhKvpwVog?e=SKfGoS>

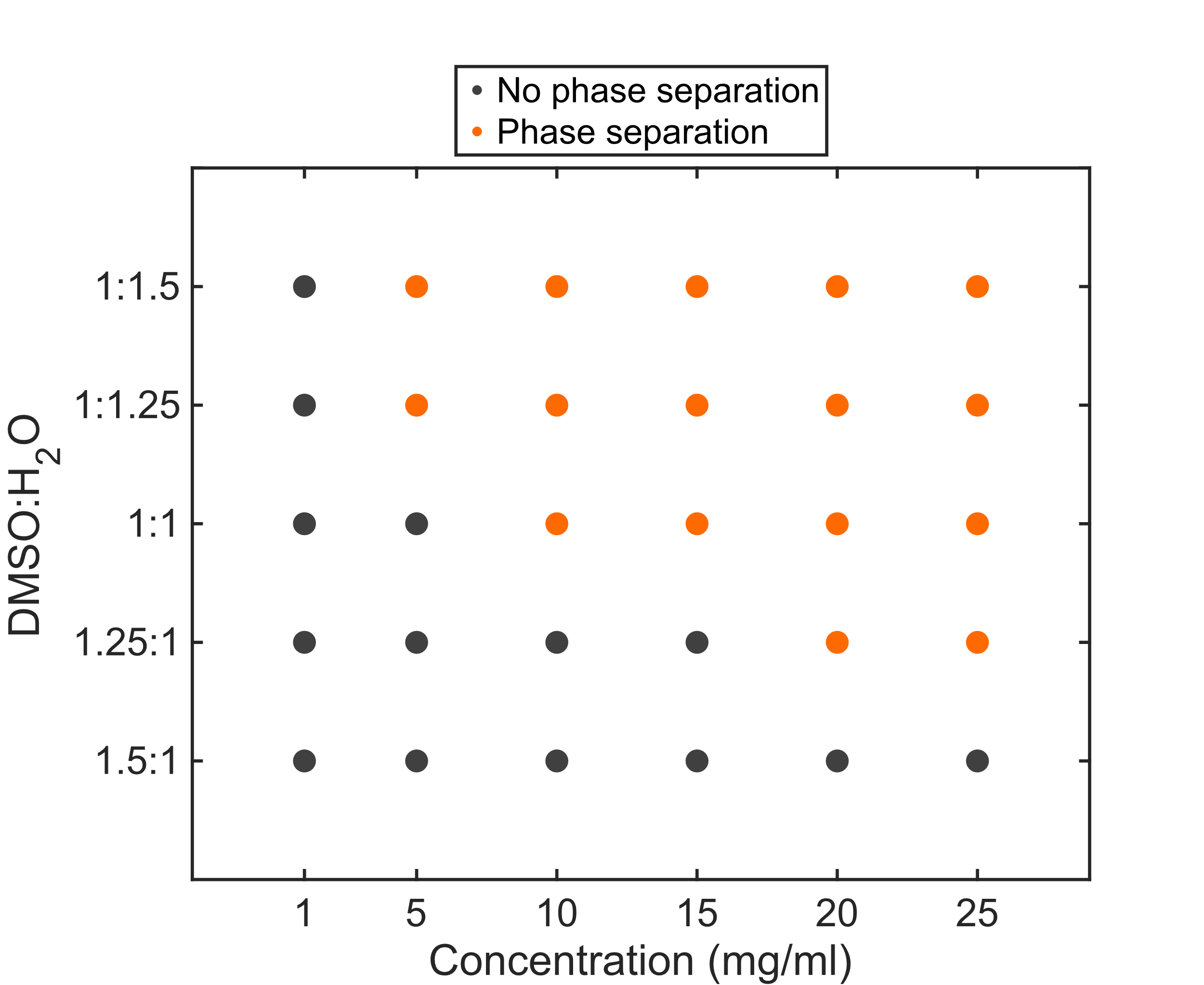

**Figure S1.** Phase diagram of z-FF prepared under different DMSO-H_2_O ratios over concentrations of 1-25 mg/ml. Data is captured after 30 minutes of aging at room temperature.

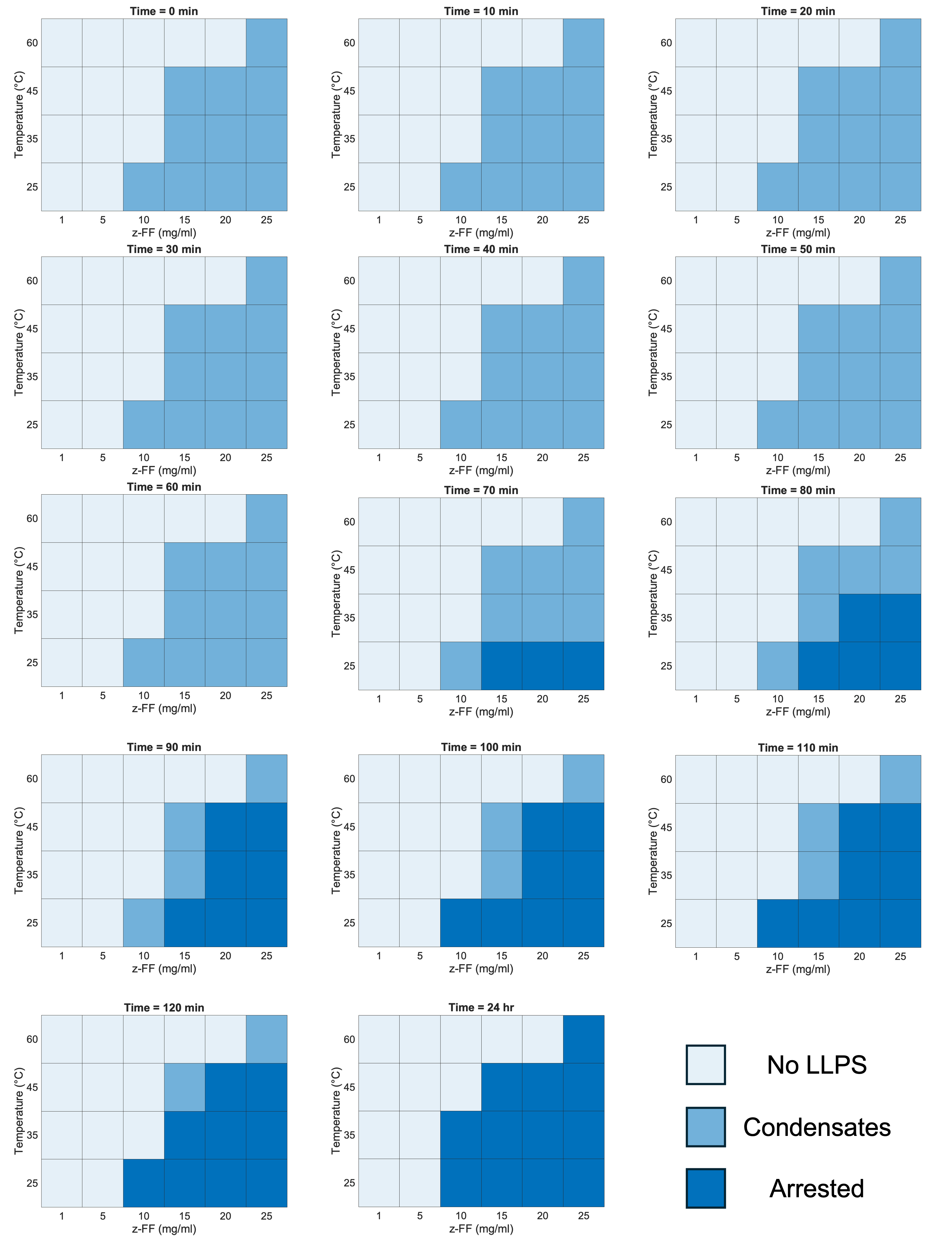

**Figure S2.** Phase diagrams of z-FF with DMSO-H_2_O ratio of 1:1 at different time points with final ThT concentrations of 100 µM.

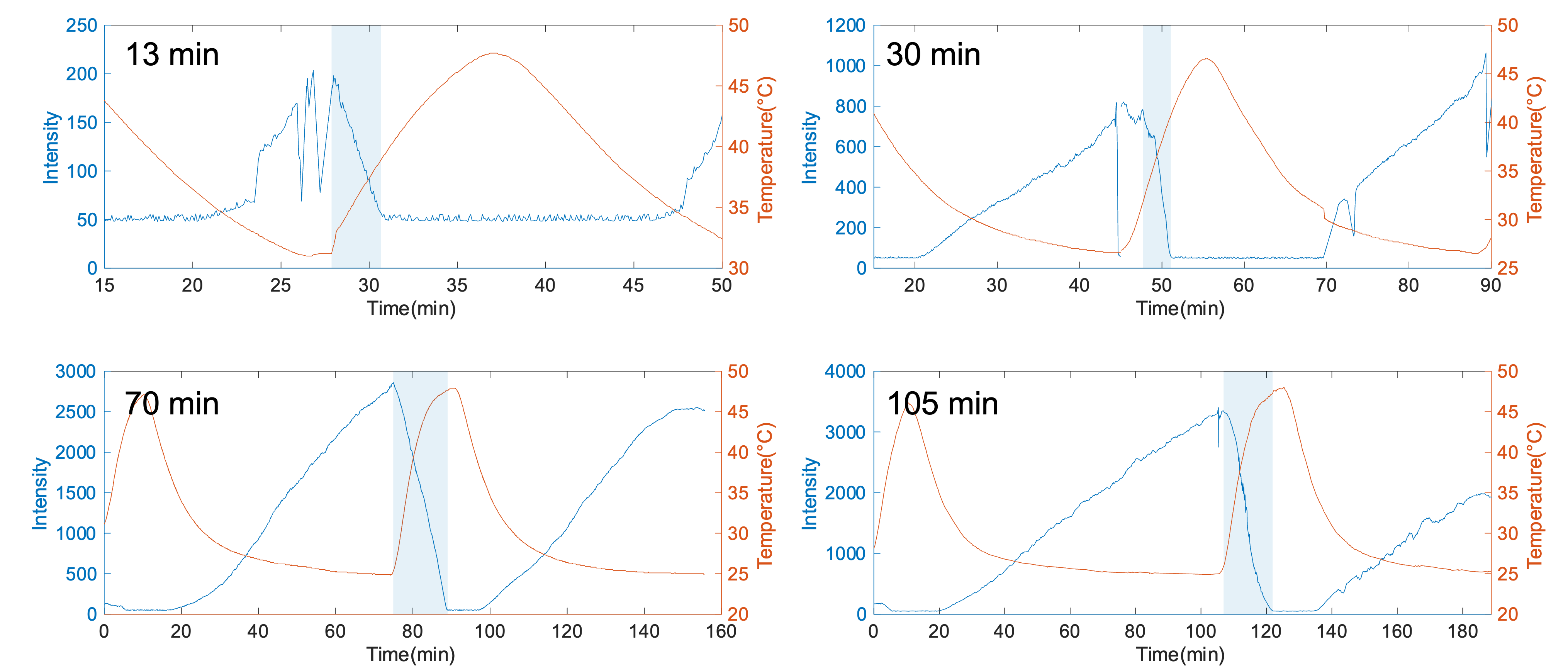

**Figure S3.** Dissolution curve for 13, 30, 70, and 105 minute-aged condensates. The areas highlighted are used to calculate the dissolution rate. The dissolution rate is calculated by the step change of normalized intensity divided by the step change of temperature. The sudden decrease in intensity at the low-temperature region is due to microscope refocusing.

The dissolution rate calculated at a given time point is calculated by:

$$Dissolution rate=\frac{\frac{dI_{norm}}{dt}}{\frac{dT}{dt}}=\frac{dI_{norm}}{dT}$$

Where $I_{norm}$ is the normalized intensity based on the region of interest, *T* the sample temperature, and *t* the time.

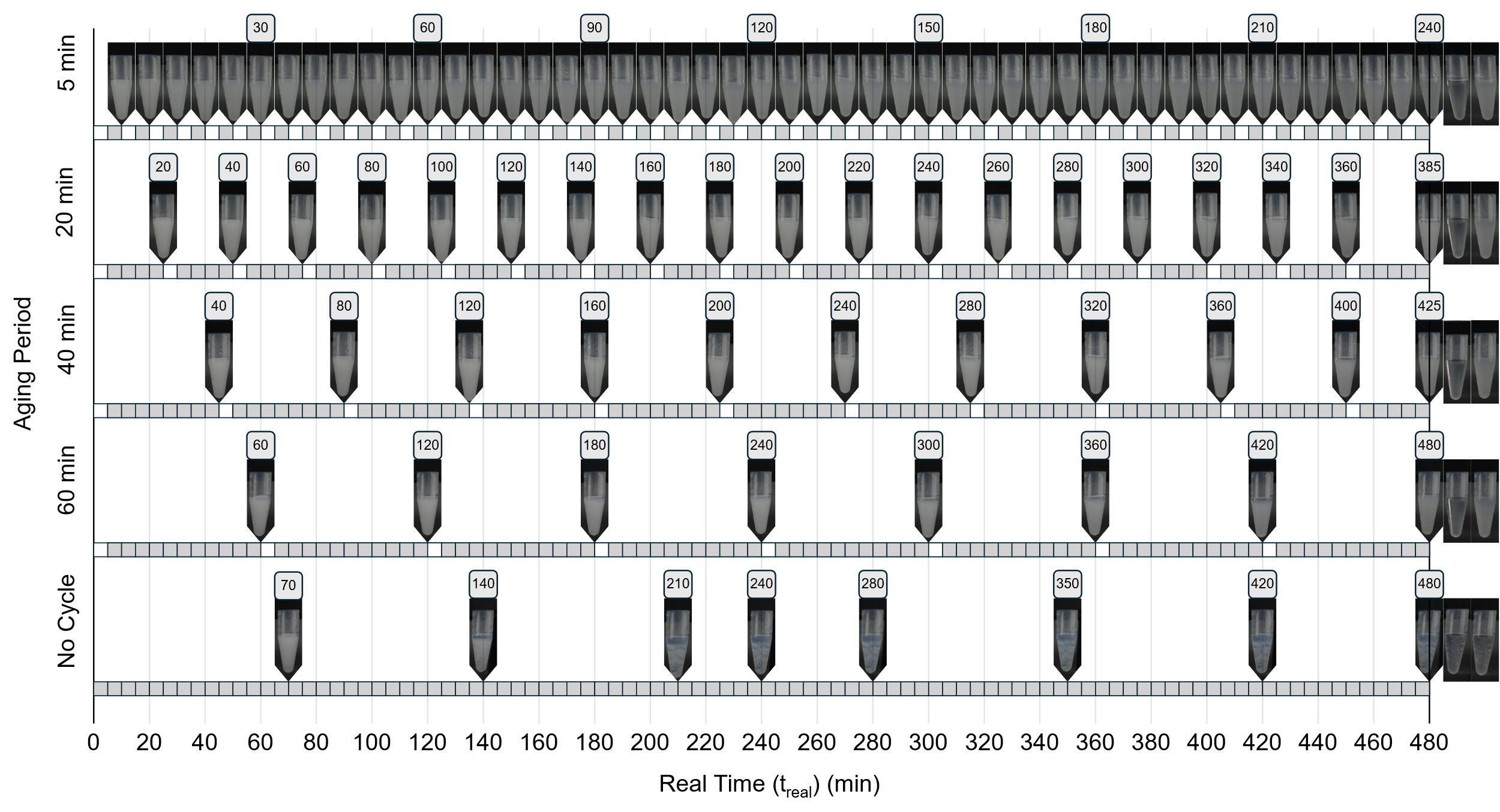

**Figure S4.** Temperature cycling experiment done in the centrifuge tubes. Each unit box corresponds to a time interval of 5 minutes with grey and white indicating the turbid and transparent phases, respectively. Grey boxes directly succeeding a white box represent the 5-minute cooling period from 65 °C to 25 °C, followed thereafter by incubation at 25 °C for each successive grey box. Both cases are included in the aging period of condensates. The cumulative aging times of samples (i.e. the age of condensates) are labelled on the top of the images. The last two images show the final heating and cooling of samples for 5 minutes to check the reversibility of condensates.

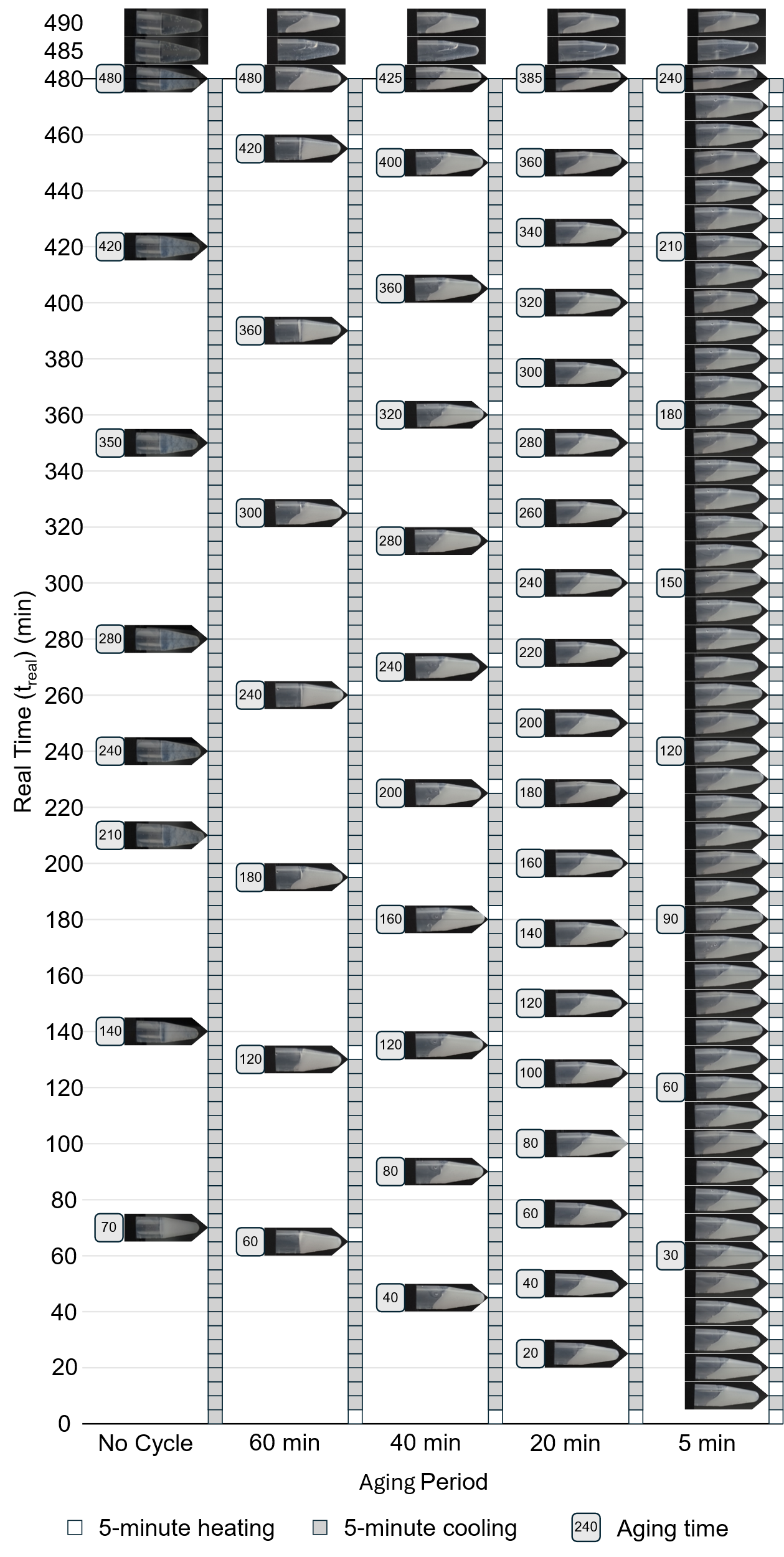

**Figure S5.** The centrifuge tube in each time point (Figure S4) is tilted to check the mobility of the liquids.

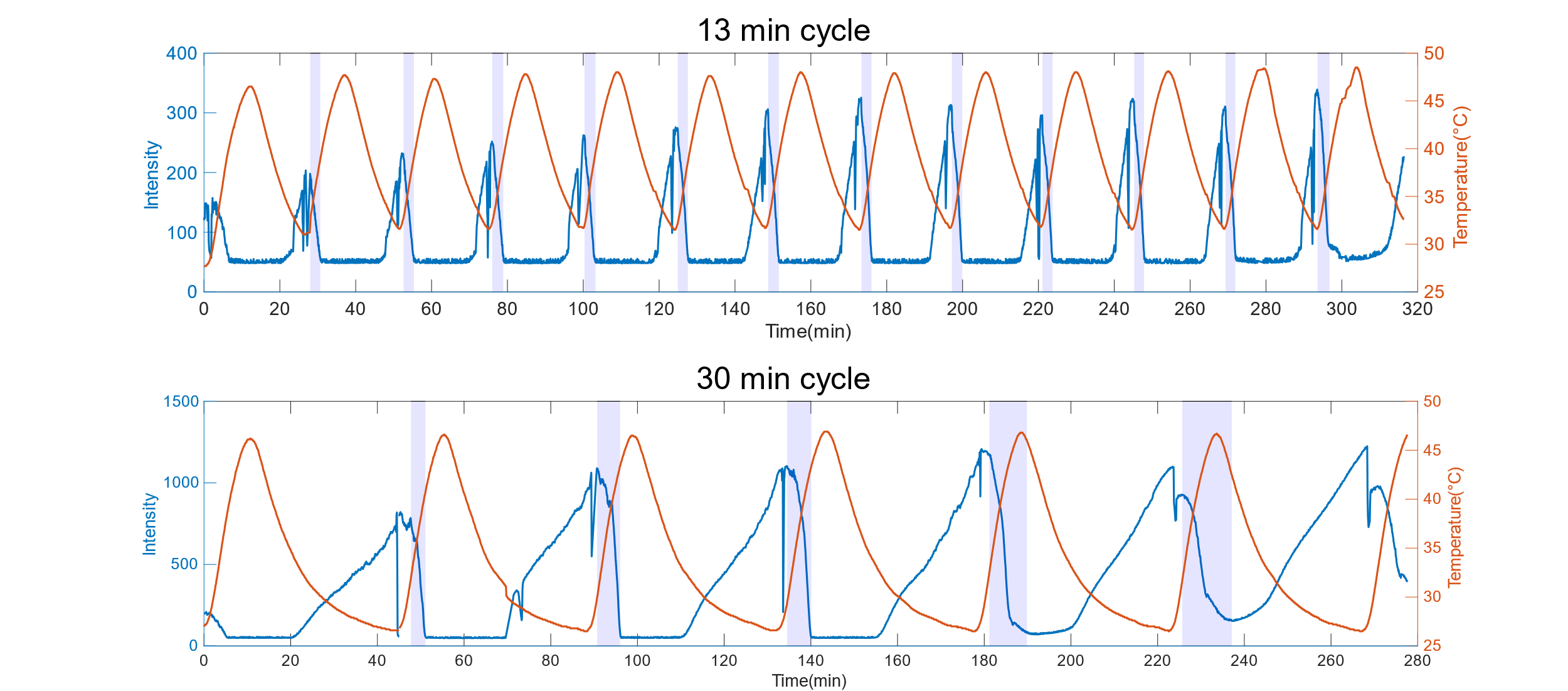

**Figure S6.** Mean intensity profiles of 13-min and 30-min cycles. The highlighted areas are used to calculate the dissolution rate using the same method mentioned in Figure S3.
